# Phosphorus uptake and toxicity is delimited by mycorrhizal symbiosis in P-sensitive *Eucalyptus marginata* but not in P-tolerant *Acacia celastrifolia*

**DOI:** 10.1101/2021.09.28.462111

**Authors:** Mark Tibbett, Matthew I. Daws, Megan H. Ryan

## Abstract

Many plant species from regions with ancient, highly-weathered nutrient-depleted soils have specialised adaptations for acquiring P and are sensitive to excess P-supply. Mycorrhizal associations may regulate P-uptake at high external P-concentrations, potentially reducing P-toxicity. We predicted that excess P-application will negatively impact species from the nutrient-depleted jarrah forest of Western Australia and that mycorrhizal inoculation will reduce P-toxicity by regulating P-uptake. For seedlings of the N_2_-fixing legume *Acacia celastrifolia* and the tree species *Eucalyptus marginata*, we measured growth at P-concentrations of 0 to 90 mg kg^−1^ soil and in relation to inoculation with the arbuscular mycorrhizal fungus (AMF) *Rhizophagus irregularis*. Non-inoculated *A*. *celastrifolia* maintained leaf P-concentrations at <2 mg g^−1^ dry mass (DM) across the range of external P-concentrations. However, for non-inoculated *E*. *marginata*, as external P-concentrations increased leaf P also increased, reaching >9 mg g^−1^ DM at 30 mg P kg^−1^ soil. *A*. *celastrifolia* DM increased with increasing external P-concentrations, while *E*. *marginata* DM was maximal at 15 mg P kg^−1^ soil, declining at higher external P concentrations. Neither DM nor leaf P of *A*. *celastrifolia* were affected by inoculation with AMF. For *E*. *marginata*, even at 90 mg P kg^−1^ soil, inoculation with AMF resulted in leaf P remaining <1 mg g^−1^ DM, and DM being maintained. These data strengthen the evidence base that AMF may not only facilitate P-uptake at low external P-concentrations, but are also important for moderating P-uptake at elevated external P-concentrations and maintaining plant P concentrations within a relatively narrow concentration range.

## 1. INTRODUCTION

Ancient and highly weathered soils, such as those found in SW Western Australia, have naturally low phosphorus (P) concentrations. Many plants adapted to growth under these low P-conditions have evolved a range of strategies for P-acquisition, including cluster roots and exudation of carboxylates and phosphatases (Lambers *et al*., 2006, 2008). A range of species also form associations with arbuscular mycorrhizal fungi (AMF), the extensive mycelial development of which expand the volume of soil from which nutrients can be scavenged (Tibbett, 2000; Tibbett & Sanders, 2002; Smith *et al*., 2015).

Species adapted to naturally low soil P concentrations may display symptoms of P-toxicity when supplied with P concentrations above those that they experience naturally in soil (Handreck, 1991; Lambers *et al*., 2002; Shane *et al*., 2004a; Standish *et al*., 2007; Pang *et al*., 2010; de Campos *et al*., 2013; Williams *et al*., 2019), due potentially to the loss of low affinity transporter systems (Huang *et al*., 2011). P-sensitive species occur in a range of families, including the Fabaceae, Haemodoraceae, Myrtaceae, Proteaceae and Rutaceae. Symptoms of P-sensitivity are highly species-specific and occur at shoot P-concentrations less than 1 mg g^−1^ dry mass (DM) to more than 40 mg P g^−1^ DM (Shane *et al*., 2004b and references therein). Symptoms of P toxicity include a reduction in growth with increasing external P (e.g., Standish *et al*., 2007; Williams *et al*., 2019) and visible symptoms including early leaf senescence and necrotic and chlorotic regions on leaves (e.g., Handreck, 1991; Lambers *et al*., 2002; Shane *et al*., 2004ab; Kariman *et al*., 2014a; Ye *et al*., 2021).

Mycorrhizal symbioses are well known for increasing P-uptake in nutrient deficient soils and increasing the P-status of host plants (Bougher *et al*., 1990; Koide & Mosse, 2004; Smith *et al*., 2011; Kariman *et al*., 2018). However, they can also enable the growth of plants in soils containing toxic concentrations of heavy metals or certain essential trace elements such as cadmium and zinc, by controlling the uptake of metal ions (Jentschke & Godbold, 2000; Hildebrandt *et al*., 2007; de Oliveira *et al*., 2020; Yazici *et al*., 2021). Similarly, AMF can modify P uptake in the host plant by reducing the expression of genes encoding high-affinity phosphate transporter proteins. While mycorrhizal associations can increase plant shoot P concentrations by increasing uptake at low P-availability and enabling exploitation of a greater soil volume (Tibbett, 2000), there is increasing evidence, at least for arbuscular mycorrhizal fungi (AMF), that they can also moderate shoot-P at high P-availability. For example, Nazeri *et al*. (2014) demonstrated, for a range of legume species, that inoculation with AMF could maintain shoot P concentrations within relatively narrow boundaries following the application of a single pulse of P. This effect was modulated by both mycorrhizal related reductions in rhizosphere carboxylates and P transport from roots to shoots.

Jarrah (*Eucalyptus marginata*) is a dominant overstorey tree in the jarrah forest of south west Western Australia and, based on pot experiments, is known to be sensitive to elevated external P (Kariman *et al*., 2014a, 2016). For example, in the absence of inoculation with AMF, Kariman *et al*. (2014a) reported the onset of leaf chlorosis and necrosis in the week following application of a pulse of P to jarrah seedlings. When seedlings were inoculated with AMF visible symptoms associated with P-toxicity and shoot-P concentrations were reduced. While studies have largely focused on visible phytotoxicity symptoms resulting from P-application (e.g., Handreck, 1991; Lambers *et al*., 2002; Shane *et al*., 2004b; Kariman *et al*., 2014a), a recent study by Williams *et al*. (2019) demonstrated that P-toxicity can also be expressed as a significant reduction in growth rates at shoot-P concentrations that do not necessarily result in visible symptoms. Consequently, there is a need to better understand both longer-term effects of applying P and potential interactions of AMF on shoot-P concentrations and plant growth.

For jarrah these observations in relation to P may also have significant practical implications: large areas of jarrah forest are cleared and restored each year following bauxite mining. Fertiliser application (especially P), to maximise early plant growth, is generally viewed as a key step in the rehabilitation process (Ward *et al*., 1990; Grant *et al*., 2007; Tibbett, 2010). However, there may be the potential for negative impacts on plant growth from applying excess P, particularly given newly restored forest have low species diversity and abundance of AMF (Gardner & Malajckuk, 1988; Glen *et al*., 2008). Our own unpublished observations have found very poor levels of colonisation by AMF (near absent) in seedlings of acacias and eucalypts in recently restored sites (Tibbett & Ryan unpublished).

In the light of potential P toxicities in tree seedlings, and the prospect for symbiotic mitigation of such effects, we investigated two linked hypotheses anticipating contrasting responses for two species with distinctive ecological strategies: *Acacia celastrifolia*, a large understorey ruderal legume that exhibits a strong positive growth as a seedling in the field and jarrah, the dominant overstorey tree which constitutes around 80% of stems in the native forest (Daws *et al*., 2015; Tibbett *et al*., 2020). Our first hypothesis was that *A*. *celastrifolia* would be highly responsive to P-addition over a wide range of exogenous supply whereas Jarrah would not and would potentially show signs of growth depression and toxicity. Based on the results of our first experiment, our second hypothesis was that AMF inoculation would alter the response in terms of P-supply, growth and uptake, leading to a suppression in acacia growth and offer remedial effect on jarrah growth response and P uptake at high amendment rates.

## 2. MATERIALS AND METHODS

### 2.1. Plant and soil material

Seeds of *Eucalyptus marginata* Donn ex Smith (Jarrah) and *Acacia celastrifolia* Benth. were collected in the northern jarrah forest of Western Australia, ca. 130 km SSE of the state capital Perth (32° 48′ S 116° 28′ E). The water impermeable seed coat of *A*. *celastrifolia* was chipped at the end furthest from the axis using a scalpel and seeds of both species were soaked for 2 h in 1:10 ‘Seed Starter’ smoke water (Kings Park Botanic Gardens and Parks Authority, Perth, WA). Seeds were sown on the surface of 10 % agar water and placed at 15°C in the dark.

For both experiments, once seeds had commenced germination, seedlings were transplanted into sealed 9.6 litre pots at a depth of 10 mm. The pots contained c. 4 litres of disinfested topsoil that had been steamed twice for three hours at 80 °C, dried at 40 °C and then sieved to 4 mm. The topsoil used in the experiment was also obtained the northern jarrah forest of Western Australia (32° 48′ S 116° 28′ E). Jarrah forest soils are gravelly with low concentrations of available N, P and K (Tibbett *et al*. 2020) and high rates of P fixation on amorphous iron and aluminium oxides. Ten germinated seeds were placed into each pot. For the second mycorrhizal inoculation experiment, spores of *Rhizophagus irregularis* (Błaszk., Wubet, Renker & Buscot) C. Walker & A. Schüßler 2010 (formerly *Glomus intraradices*) were placed at the bottom of the hole before the seeds were added. Pots were watered approximately weekly to 50 % field capacity and seedlings were thinned to the two healthiest plants after 21 days, and then down to one plant after 31 days.

To ensure that only P was limiting in the experiment, and that no nutrient imbalances were induced by the addition of P, 10 ml of modified Long Ashton’s nutrient solution (minus P) was added to each pot 15 days after seedlings were planted (Cavagnaro *et al*., 2001). Macronutrients: K_2_SO_4_ (20 mM), MgSO_4_·7H_2_O (15 mM), CaCl_2_·2H_2_O (30 mM), FeEDTA (1 mM), (NH_4_)2SO_4_ (40 mM), NaNO_3_(80 mM). Micronutrients: H_3_BO_3_ (28.6 mg l^−1^), MnCl_2_·4H_2_O (18.1 mg l^−1^), ZnSO_4_·7H_2_O (2.2 mg l^−1^), CuSO_4_·5H_2_O (0.8 mg l^−1^), NaMoO_4_·2H_2_O (0.25 mg l^−1^).

### 2.2. Experimental design

Twenty-three days after planting, phosphate was added to the P-treatments in the form of potassium dihydrogen phosphate (KH_2_PO_4_). To ensure a constant ionic background and balanced potassium levels, potassium chloride (KCl) was added in inverse proportions to KH_2_PO_4_ amendments. In **experiment 1**, eight rates of P-application were used (equivalent to 0, 0.9, 4.5, 13.5. 22.5, 31.5 40.5 and 81 mg elemental P kg^−1^ soil). In **experiment 2** there were four P-application rates (equivalent to 0, 4.5, 30 and 90 P kg^−1^ soil) in a two-way factorial combination of P-application rate × mycorrhizal treatment. Both experiments were established in randomised blocks in a glasshouse, with each block containing all treatments. The temperature-controlled glasshouse was maintained at temperatures between 18°C and 28°C. Pots were regularly re-randomised throughout the growing period. All treatments were replicated six times for experiment 1 and four times for experiment 2.

### 2.3. Mycorrhizal activity (experiment 2)

Thirteen days after planting and at the end of the experiment (188 days), seedlings were screened for evidence of colonisation by AMF. Roots were cleared using KOH then stained using lactic-glycerol blue and examined by light microscopy for evidence of colonisation (Brundrett *et al*., 1996). At the end of the experiment, AM spores were also extracted from 150 g of soil sampled away from the middle of the pots following wet sieving and sucrose centrifugation (Walker *et al*., 1982).

### 2.4. Plant measurements

For **experiment 1**, plants were harvested 213 days after sowing and roots and shoots dried separately at 70 °C for dry weight determination. For **experiment 2**, plants were harvested for dry mass (DM) determination 53 or 188 days after sowing. In both experiments, plants were carefully removed from the growing medium, roots washed with water and the plants separated into roots and shoots.

### 2.5. Foliar phosphorus concentrations

Dried leaf material from the 213-day (experiment 1) and 188-day harvest (experiment 2) was ground and then subsampled for digestion. Leaf material was digested using a HNO_3_/HClO_4_ mixture with the diluted digest approximately 10% V/V with respect to HClO_4_ (70% W/W). Phosphorus content was determined using the molybdovanadophosphate method (yellow) and a spectrometer reading at a wavelength of 460 nm (modified from Simmons, 1975 and 1978).

### 2.6. Statistical analysis

For experiment one, two-way ANOVA implemented in Minitab 14 was used to assess, whether there were effects of (1) species and (2) increasing external P concentration, on either plant DM or leaf P concentration. Data did not require transformation before analyses as the assumptions of ANOVA with respect to normality and homogeneity of variances were met. For experiment 2, two-way ANOVA was used to assess, for each of the two species, the effect of (1) inoculation with AMF and (2) external P concentration on either plant DM or leaf P concentration. Finally, for experiment 2, for the plants of each of the study species inoculated with AMF, two-way ANOVA was used to assess the interaction of (1) study species and (2) external P concentration on spore count in the soil at the end of the experiment. Spore counts were log_10_(*n* + 1) transformed to ensure normality.

## 3. RESULTS

### 3.1. Effect of external P concentration on dry mass and leaf P of non-mycorrhizal plants (experiment 1)

In experiment 1 with non-mycorrhizal plants, there was a significant effect of P-application rate on plant DM after 213 days (Two-way ANOVA *P* < 0.001), but the two species (*A*. *celastrifolia* and *E*. *marginata*) responded differently to applied-P (Two-way ANOVA *P* < 0.05; Figure 1). For *A*. *celastrifolia* there was an initial rapid increase in DM as P-application rate increased from 0 to 15 mg kg^−1^ soil. At higher P-application rates, the rate of increase in DM declined. Nonetheless, total DM at the highest application rate (81 mg kg^−1^ soil) was 10× higher than at 0 mg P kg^−1^ soil (Figure 1). For *E*. *marginata*, there was also an initial increase in DM as P-application rate increased from 0 to 15 mg kg^−1^ soil (Figure 1). However, at P-application rates greater than 15 mg kg^−1^, DM declined: plant mass at the P-application rate of 15 mg kg^−1^ was more than twice that at 81 mg kg^−1^ (Figure 1).

**Figure 1.**
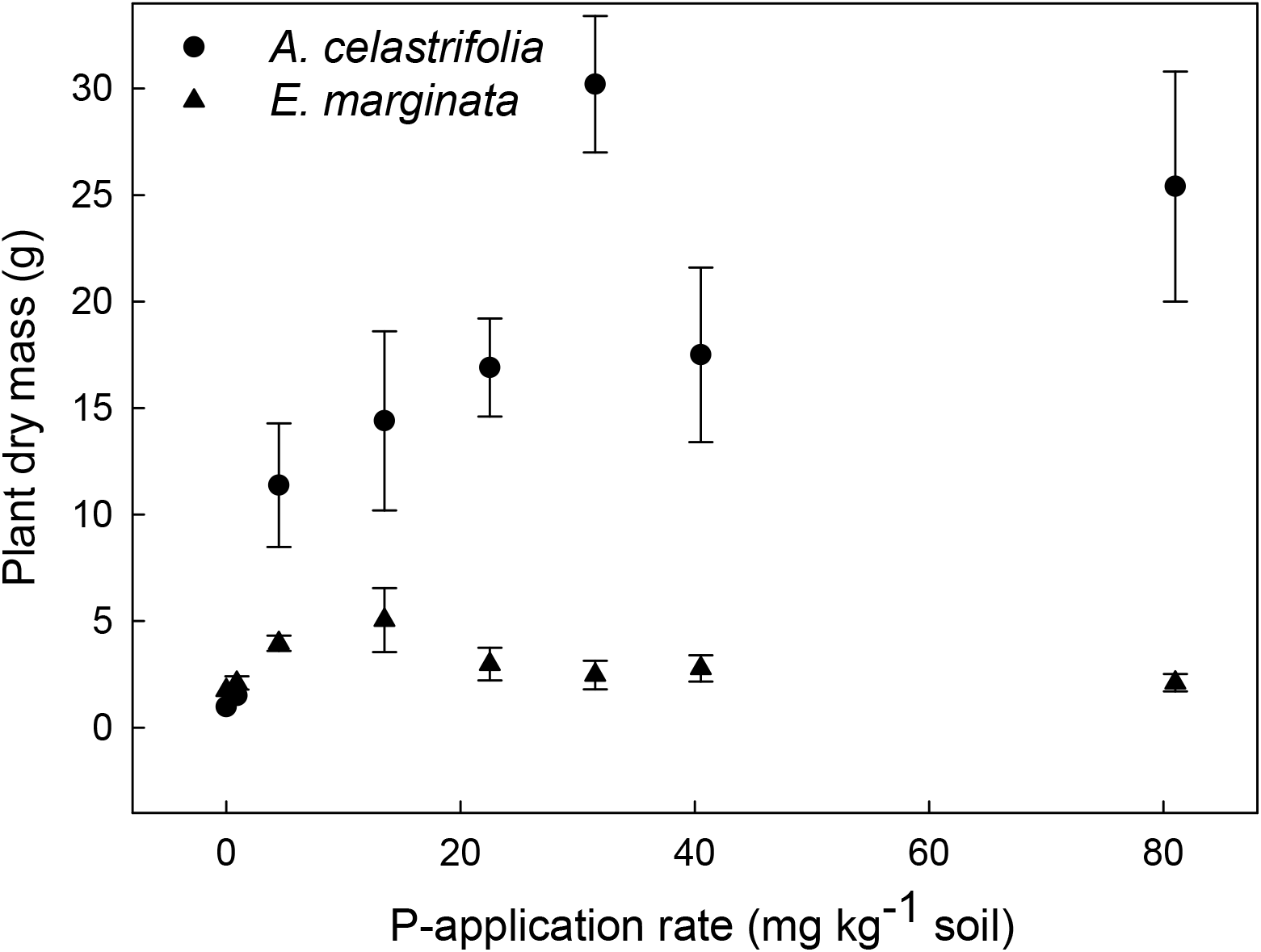
The effect of P-application rate on plant dry mass of non-mycorrhizal *Acacia celastrifolia* and *Eucalyptus marginata* assessed 213 days after sowing. Bars ± 1 SE.

In experiment 1, leaf P concentrations of *A*. *celastrifolia* were ca. 0.65 mg g^−1^ DM for plants at the nil P application-rate, then increasing to a maximum of ca. 3.2 mg g^−1^ DM at the P-application rate of 81 mg kg^−1^ soil (Figure 2). For *E*. *marginata* leaf P was ca. 0.22 mg g^−1^ DM at nil P-applicator rate (Figure 2) and increased rapidly with P application reaching ca. 9 mg g^−1^ DM at the P-application rate of 30 mg kg^−1^. As the P- application rate increased to 80 mg kg^−1^, leaf P concentrations remained at ca. 9 mg g DM (Figure 2). These responses were reflected in significant main effects of P-application rate and species as well as a significant P-application × species interaction (Two-way ANOVA, *P* < 0.001), a significant (*P* < 0.001) and a non-significant main effect of P-application (*P* < 0.001) on leaf P. For *E*. *marginata* the leaf P concentration corresponding to maximum plant DM accumulation, and above which DM accumulation declined, was ca. 4 mg g^−1^ DM (Figures 1 and 2).

**Figure 2.**
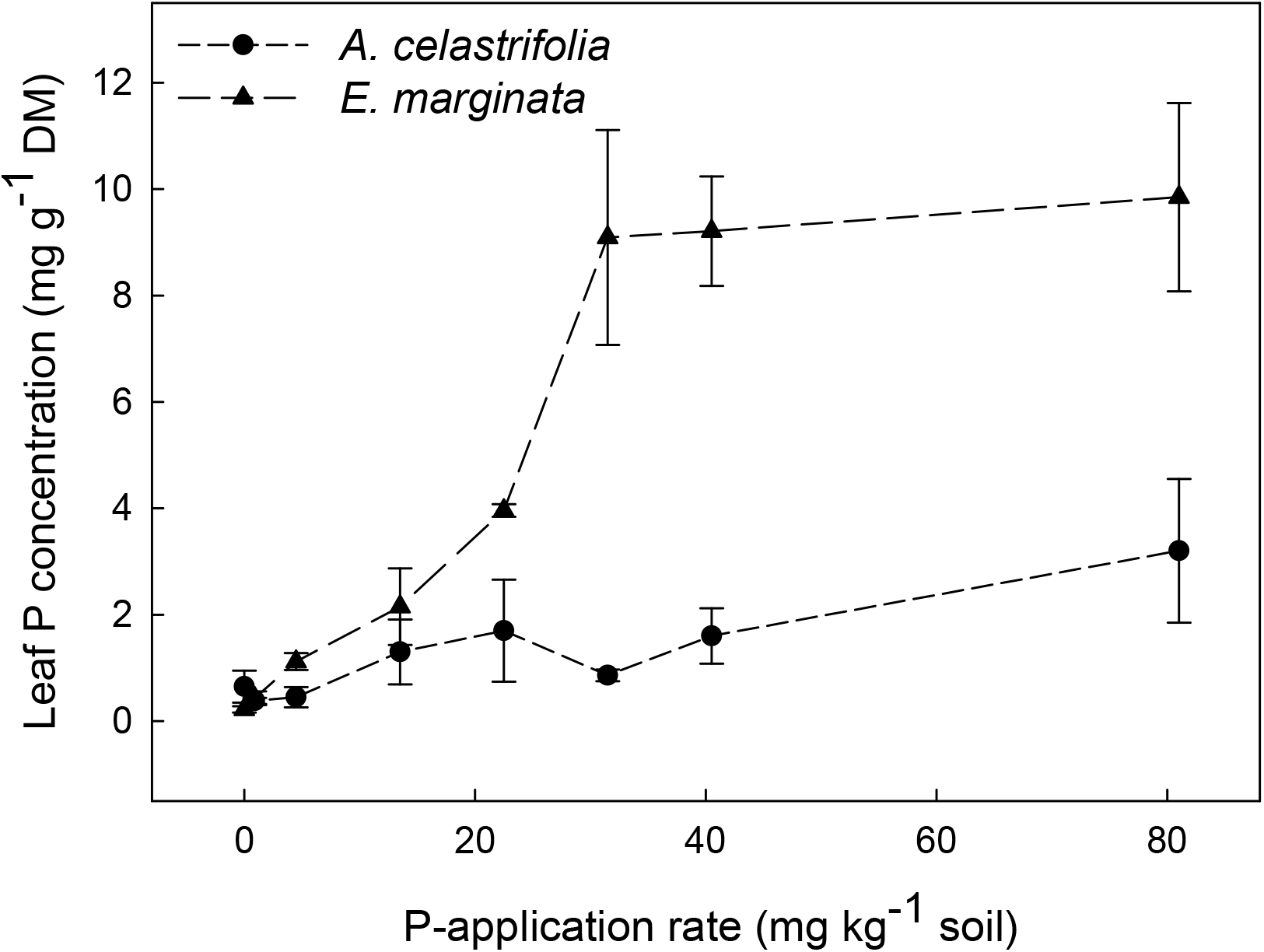
The effect of P-application rate on leaf P-concentration for *Acacia celastrifolia* and *Eucalyptus marginata* assessed 213 days after planting. Bars ± 1 SE.

### 3.2. Effect of AM colonisation on dry mass and leaf P (experiment 2)

For experiment 2, evidence of early colonisation (6-29 %) by AM was found in four of the twelve samples taken (one *A*. *celastrifolia* and three *E*. *marginata*). Roots taken at the end of the experiment did not clear properly and were not able to be assessed reliably and consequently were discarded. Spore counts from mycorrhizal pots were low (under 100 per 150 g soil) for both *A*. *celastrifolia* and *E*. *marginata* at the nil P-application rate. There was a significant effect of plant species on sporulation in the inoculated treatment (Two-way ANOVA, *P* < 0.01) with sporulation varying greatly between the species at the nil P-application rate, peaking at 4.5 mg kg^−1^ P for *E*. *marginata* yet remaining fairly low at all P application rates (Table 1). In contrast, sporulation continued increased with P application for *A*. *celastrifolia*, peaking at 90 mg kg^−1^ P with a mean of 276 spores per 150 g of soil. This difference in response between the two species was reflected in the P application rate × species interaction being highly significant (Two-way ANOVA, *P* < 0.001), but the main effect of P application being non-significant (*P* > 0.05).

**Table 1.** The effect of plant species and phosphorus (P)-application rate on the spore count of the arbuscular mycorrhizal fungus *Rhizophagus irregularis*. Spore counts were taken at the end of experiment 2 (day 188). Error bars are ± 1 SE of the mean.

| Plant species | P-application rate<br>(mg kg<sup>-1</sup> soil) | Spore count |
| --- | --- | --- |
| *Acacia celastrifolia* | 0 | 45.3 ± 4.4 |
| | 4.5 | 117.3 ± 13.0 |
| | 30 | 191.8 ± 27.0 |
| | 90 | 276.0 ± 52.5 |
| *Eucalyptus marginata* | 0 | 80.5 ± 3.5 |
| | 4.5 | 103.3 ± 9.0 |
| | 30 | 58.5 ± 5.8 |
| | 90 | 37.0 ± 4.3 |

In experiment 2 with both plants inoculated and non-inoculated with AMF, for the two time periods that were measured (53 and 188 days), there was a significant effect of P-application rate on DM (Figure 3AB; Two-way ANOVA, *P* < 0.01). *A*. *celastrifolia* DM increased with P-application rate and the rate of increase declined above a P-application rate of 30 mg kg^−1^ soil. For *A*. *celastrifolia*, there was no effect of AM inoculation on DM accumulation for either measurement interval (Figure 3AB; Two-way ANOVA, *P* > 0.05).

**Figure 3.**
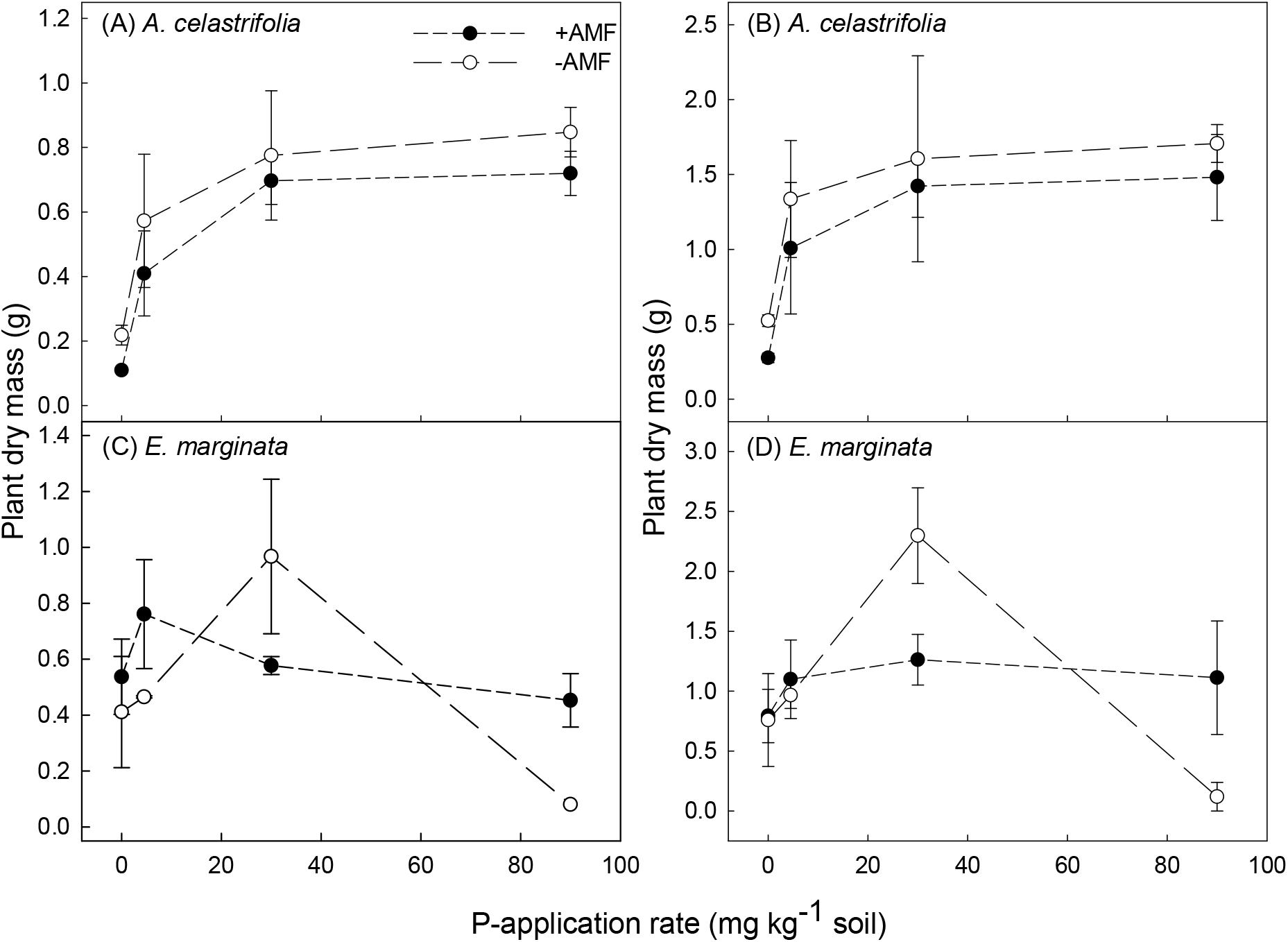
The effect of P-application rate and inoculation with arbuscular mycorrhizal fungi (AMF) on plant dry mass assessed either 53 (A and C) or 188 days after planting (B and D) for *Acacia celastrifolia* and *Eucalyptus marginata*. Note that due to elevated mortality at high P, the 53-day data point for *E*. *marginata* at 90 mg P kg^−1^ consists of data from one plant only. Bars ± 1SE.

At both measurement intervals, DM of non-inoculated *E*. *marginata* plants increased reaching a maximum at the P-application rate of 30 mg kg^−1^ soil and declined thereafter (Figure 3CD). At a P-application rate of 90 mg kg^−1^ soil, there were visible symptoms of P-toxicity including leaf necrosis. Further, for the replicate sampled at 53 days after planting, only one of the four replicate plants was still alive. For plants inoculated with AMF, there was no effect of P-application rate on DM accumulation. Further, neither a decline in DM nor visible symptoms of P-toxicity were observed at the highest P-application rate of 90 mg P kg^−1^ soil (Figure 3CD). At the second measuring interval these responses were reflected in a significant P-application rate × AMF inoculation interaction (Two-way ANOVA, *P* < 0.001). Profound differences between inoculated and non-inoculated plants can be seen in Figure 4.

**Figure 4.**
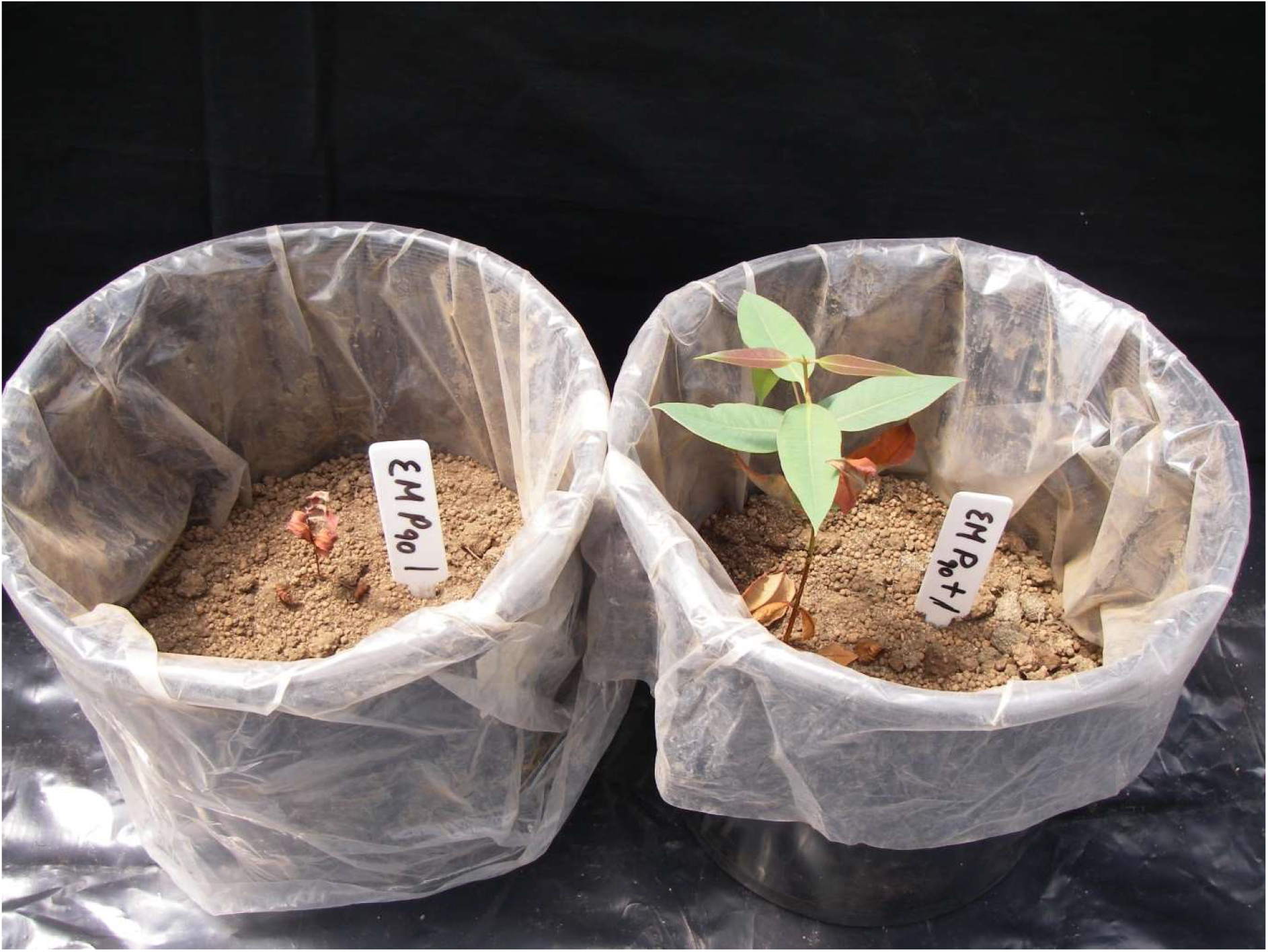
Photograph of jarrah (*Eucalyptus marginata*) seedlings from **experiment 2** (see methods section) grown at a P-application rate of 90 mg kg^−1^ soil. The pot on the left was not inoculated with spores of arbuscular mycorrhizal fungi whereas the pot on the right was inoculated with spores of AMF at the commencement of the experiment. The photograph was taken at the end of the experiment (day 188 after planting).

Leaf P concentration of *A*. *celastrifolia* increased from ca. 0.6 to 2 mg g^−1^ DM as the P-application rate increased from 0 to 90 mg kg^−1^ soil. This response was independent of inoculation with AMF (Figure 5A; Two-way ANOVA, *P* > 0.05). For non-inoculated *E*. *marginata* plants, the leaf P concentration increased from ca. 0.2 to 9 mg g^−1^ DM as the P-application rate increased from 0 to 90 mg kg soil^−1^ (Figure 5B). However, for plants of *E*. *marginata* inoculated with AMF, leaf P concentration did not respond to an increasing P-application rate and remained at ca. 0.5 mg g^−1^ DM across the entire range of P-applications rate which was reflected in a significant P application rate × AM inoculation interaction term (Two-way ANOVA, *P* < 0.001; Figure 5B).

**Figure 5.**
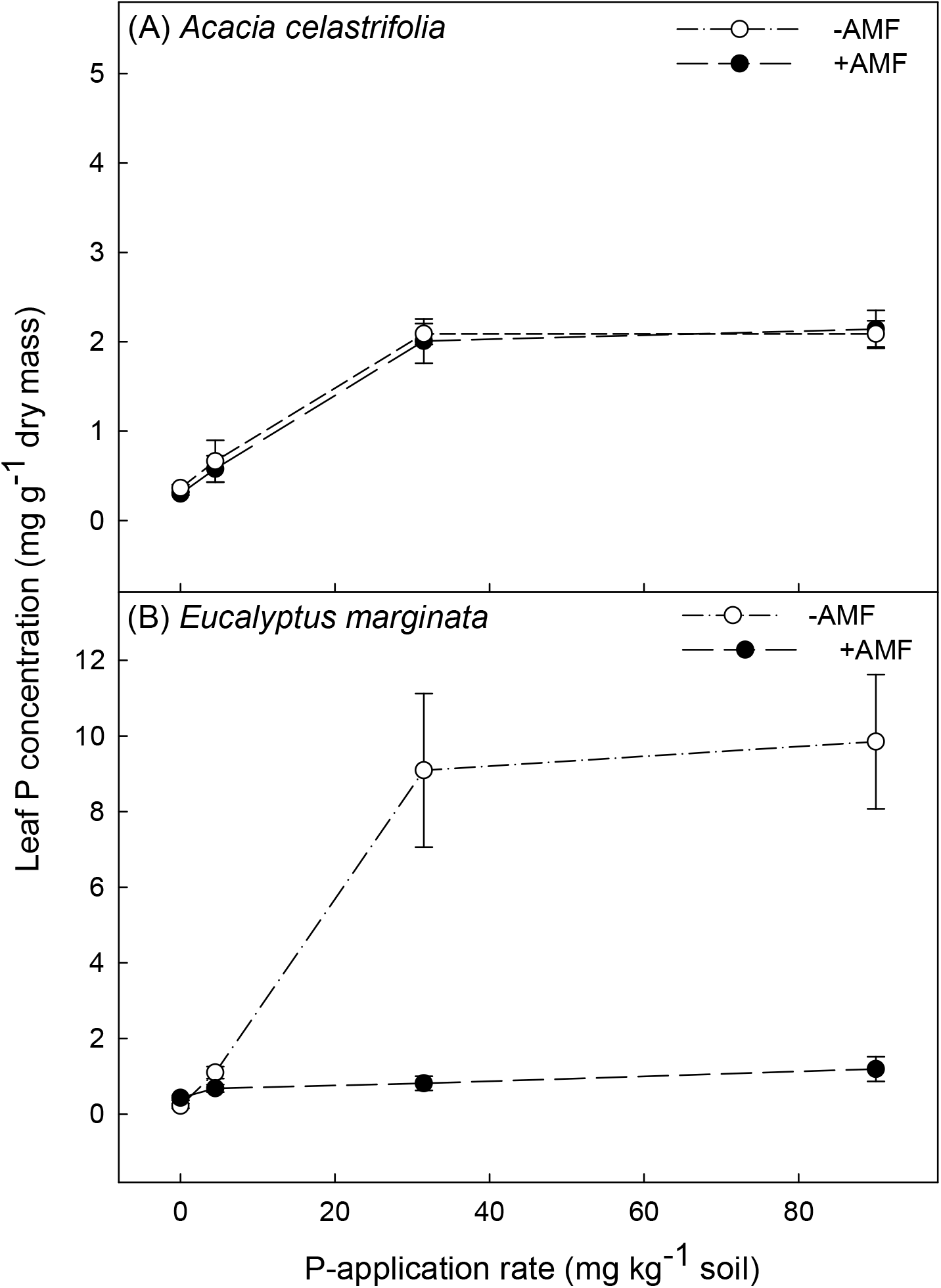
The effect of P-application rate and inoculation with arbuscular mycorrhizal fungi (AMF) on leaf P-concentration for *Acacia celastrifolia* and *Eucalyptus marginata* assessed 188 days after planting. Bars ± 1 SE.

## 4. DISCUSSION

For non-mycorrhizal plants of *A*. *celastrifolia*, an increasing P-application rate increased DM up to high level application rates (80 mg P kg^−1^ soil). However, for non-mycorrhizal plants of *E*. *marginata*, an increasing P-supply increased growth only at relatively low external P concentrations: thereafter DM declined with increasing P supply. Similar contrasting patterns in response to increasing P-supply have been observed previously for a range of Australian species from severely nutrient-impoverished environments (Grundon, 1972; Groves & Keraitis 1976; Handreck, 1997; Pang *et al*. 2010; de Campos *et al*., 2013, Williams *et al*., 2019). For *E*. *marginata*, but not *A*. *celastrifolia*, inoculation with AMF reduced growth at moderate P supply but facilitated growth at high P supply by regulating leaf P concentrations.

For non-mycorrhizal *E*. *marginata*, maximum DM was observed at P-application rates of 15-30 mg kg^−1^ soil, before declining at higher P-application rates. Similarly, maximum DM of a range of Australian natives has been reported to occur across a similar range of P supply (e.g., Bougher *et al*., 1990; Ryan *et al*., 2009; Pang *et al*., 2010; Williams *et al*., 2019). For example, for the 11 species studied by Pang *et al*. (2010), maximum growth occurred at P-application rates in the range 12-24 mg P kg^−1^. Further, for 8 of the 11 species, growth declined at P-application rates greater than those required for maximum plant DM. Shane *et al*. (2004b) reported that, for a range of species, P-toxicity occurred at leaf P concentrations of 0.9 to 47 mg g^−1^ DM. The leaf P concentrations at which we observed negative effects on growth of *E*. *marginata* (> 4 mg g^−1^ DM) are at the lower end of these reported values. However, Williams *et al*. (2019) reported for *Eucalyptus torquata* that a reduction in growth occurred at leaf P concentrations > 2 mg g^−1^ DM. One possible explanation for the difference between our current values and those reported by Shane *et al*. (2004b) is that the values reported by Shane and co-workers are for the onset of *visible* symptoms of P-toxicity (e.g., necrosis): our results and those of Williams *et al*. (2019) indicate the onset of a negative effect on plant growth and not necessarily the onset of visible symptoms.

Leaf P concentrations of non-mycorrhizal *A*. *celastrifolia* initially increased with increasing P supply. However, even at higher P supply, leaf P concentrations did not exceed ca. 2 mg g^−1^ DM. Similarly, Williams *et al*. (2019) reported that *Acacia acuminata* exhibited an initially increasing shoot P concentration in response to increasing P supply, but as P application rates increased further, shoot P concentration was maintained at ca. 2 mg g^−1^ DM. Conversely, *Acacia hemiteles* exhibited increasing shoot P with increasing P supply: shoot P reached concentrations of ca. 8 mg g^−1^ DM, and growth reduced as shoot P concentration continued to increase (Williams *et al*., 2019). de Campos *et al*. (2013) also reported on the ability of two *Acacia* species (*Acacia truncata* and *Acacia xanthina*) to regulate internal P-concentrations in relation to external P-concentrations. *Acacia truncata* was unresponsive in terms of DM accumulation as the external P concentration increased, whilst *Acacia xanthina* exhibited declining DM as P increased. These results suggest that the growth response to elevated P, even within co-occurring members of a genus, can be unpredictable as also reported for the genus *Banksia* (de Campos *et al*., 2013).

We found limited evidence of colonisation by AMF in either species with evidence of colonisation present in just four of the twelve samples examined. Similarly, low levels of colonisation have been reported in previous studies with jarrah forest species. For example, Kariman *et al*. (2012) reported that although evidence of colonisation by AMF of *E*. *marginata* in a pot experiment was found in as few as 2.3% of samples, beneficial effects of AMF were still observed. Further, it should be noted that root colonisation is not necessarily required for positive physiological responses in plant–fungus interactions (Neumann, 1959; Kariman *et al*., 2014b). Similar results have also been reported for ectomycorrhizal fungus colonisation on seedlings of *Eucalyptus diversicolor* where in the absence of applied P, despite colonisation rates on roots ranging from only 1 to 6%, there was still a significant growth benefit for seedlings resulting from inoculation (Bougher *et al*. 1990).

For *A*. *celastrifolia*, plant DM was consistently, but not significantly, lower in the AM inoculation treatment across the entire range of P-application rates. While AMF can increase P-uptake and result in increased growth (Smith *et al*. 2015), the transfer of carbon from the host plant to the AMF can also result in negative effects on plant growth. In comparison, inoculation with AMF had three contrasting effects on DM of *E*. *marginata*.

First, at a low P-application rate (4.5 mg P kg^−1^), at 53 days after planting, there was a positive classical growth effect of inoculation on DM. While inoculation with AMF did not increase leaf P concentrations it had increased total P uptake. Similarly, Kariman *et al*. (2014a) reported for 14-week-old *E*. *marginata* seedlings grown under P-deficient conditions that inoculation with the AMF *R*. *irregularis* did not increase leaf P concentration. In contrast, at low external P concentrations Jones *et al*. (1998) reported that inoculation with AMF both increased shoot P concentrations and growth of *Eucalypus coccifera*.

Second, at P-application rate, inoculation with AMF significantly depressed growth suggesting that at this supply, the association was parasitic rather than mutualistic. Indeed, for a range of species at elevated P concentrations, associations with AMF have been shown to move from mutualistic to parasitic (Johnson *et al*., 1997; Hoeksema *et al*., 2010; Johnson, 2010).

Third, at high P-application rates, inoculation with AMF significantly increased DM, compared to non-inoculated plants, whilst maintaining leaf P concentrations within a similar range to that observed at lower external P concentrations. This is not a classical plant growth effect, rather a suppression to toxicity due to the symbiosis. We posit this mechanism is related to a (down)regulation of root epidermal transporters which has been observed in AM plants for P, and at high concentrations for cadmium and putatively arsenic (De Oliveira *et al*., 2020; Kariman *et al*., 2014a; Kariman *et al*., 2016; Nazeri *et al*., 2014). In nature, where plants are commonly mycorrhizal (Kariman *et al*., 2018) this may be a common mechanism whereby plants are protected from toxicities (at least to some extent) by mycorrhizal symbiosis.

Except for our control, nil P treatment, leaf P concentrations for both of our study species when non-mycorrhizal were higher than concentrations previously reported in plants growing in relatively undisturbed and unfertilised jarrah forest. For example, values of 0.4-0.45 mg P g^−1^ DM have been reported for *E*. *marginata* (Hingston *et al*., 1981, M.I. Daws unpublished data) and 0.3 mg P g^−1^ DM for *A*. *celastrifolia* (M.I. Daws unpublished data). For *E*. *marginata* inoculated with AMF, leaf P was maintained at concentrations similar to those observed in unfertilised forests across the full range of P-application rates. Since P is widely applied to newly established *E*. *marginata* stands following post mining rehabilitation (Standish *et al.* 2015) investigating the potential role of colonisation by AMF in moderating P-uptake in the field would be of value, particularly since colonisation of roots in newly established sites may be limited by the availability of propagules, e.g., spores, hyphae, colonised roots (Jasper *et al*., 1991).

AMF are generally viewed as being important for increasing P-uptake and facilitating growth at low external P-concentrations. However, our data support a growing understanding that that by regulating plant P concentration within a sufficient concentration range, AMF play an important role at high external P concentrations in enabling plant growth at concentrations that would otherwise result in reduced growth and P-toxicity.

## AUTHOR CONTRIBUTIONS

MT and MR conceived the study. MT designed the study. MD analysed the data. MT and MD interpreted the data and wrote the manuscript. All authors contributed to the draft manuscript.

## ACKNOWLEDGEMENTS

Our thanks to Russell Beazley, Anna Dudley, Matt Braimbridge, Henning Wallrabenstein and Bridget Kennedy for their contribution to this work.

## DATA AVAILABILITY STATEMENT

Data sharing is not applicable to this article as all new created data is already contained within this article.

